# Glomerular immune injury promotes anti-tumor activity

**DOI:** 10.1101/2023.10.14.562333

**Authors:** Shimrit Avraham, Ben Korin, Jerry Hung-Hao Lo, Mayra Cruz-Tleugabulova, Hari Menon, Spyros Darmanis, Yuxin Liang, Zora Modrusan, Steffen Durinck, Joshua D Webster, Andrey S Shaw

## Abstract

Recent evidence suggests that the interaction between the tumor microenvironment (TME) and systemic host environment can alter the host immune system to promote anti-tumor activity. Here, we investigated whether glomerular immune injury affects cancer progression. We used nephrotoxic serum nephritis (NTN), a model for glomerular immune injury, and followed it by cancer cell implantation. NTS-injected mice developed smaller primary tumors compared with controls. Tumors of NTS-injected mice had more activated CD8 T cells, suggesting a role for the immune system in the anti-tumor phenotype. Using RNA-seq data, we identified transcriptomic alterations in the bone marrow following NTN. Moreover, using scRNA-seq of white blood cells following NTN we found these transcriptomic alterations were reflected in γδ T cells and neutrophils. This is the first study to show that glomerular immune injury changes the transcription of cells in the bone marrow to advance anti-tumor activity. Our study highlights the pivotal role of BM-mediated transcriptional alterations underlying the enhanced host immunity to tumor growth.

## Introduction

It has been well documented that inflammatory cell infiltration into the tumor microenvironment (TME) affects tumor fate^1,2^. The host immune system has suppressive effects on tumor progression. Cytotoxic T cells and NK cells can recognize and eliminate tumor cells, thereby suppressing tumor growth. Moreover, the presence of immune cells within the TME is associated with improved prognosis and better patient survival^3–6^. On the other hand, host immune cells, such as tumor-associated macrophages (TAMs) and myeloid-derived suppressor cells (MDSCs), can promote tumor growth by releasing growth factors and cytokines that stimulate angiogenesis and cell proliferation. Additionally, these immune cells can create an immunosuppressive environment by inhibiting the activity of cytotoxic T cells and natural killer (NK) cells^7,8^. Alternatively, they can play a dual role, for example, tumor-associated neutrophils (TANs) can be antitumorigenic or pro-tumorigenic^9^. Circulating factors from distant injured tissues have been shown to affect the TME^10,11^. Furthermore, the interplay between the tumor microenvironment and systemic host milieu can alter the host immune system to either promote or inhibit cancer progression^12–14^. In this context, it was recently established that immune events could induce a trained phenotype, producing more vigorous immune responses to secondary stimuli^14–17^. While cancer outcomes following pathogenic exposure have been previously described, the effect of glomerular immune injury on tumor growth is unknown.

Here, we studied whether glomerular immune injury affects cancer progression and spread. For that, we used nephrotoxic serum nephritis (NTN), an experimental model for immune complex-mediated glomerulonephritis. Subsequent to glomerular injury, we implanted mice with two different cancer models, a lung cancer model (Lewis lung carcinoma [LL/2]) and a colon adenocarcinoma model (MC38). We found that mice that were injected with nephrotoxic serum (NTS) developed smaller primary tumors than those implanted in control mice, indicating that glomerular immune injury attenuates tumor growth. Moreover, by analyzing immune cells in the bone marrow (MB) and blood of non-tumor-bearing control and NTS-injected mice, we found that NTN induces transcriptomic changes in BMCs and blood neutrophils. These findings suggest that BM-mediated transcriptional alterations underlie the enhanced host immunity to tumor growth.

## Methods

### Ethical statement

All animal procedures were conducted under protocols approved by the institutional AAALAC-accredited review board and were performed following the Guide for the Care and Use of Laboratory Animals.

### Animal models

Age-matched C57BL/6J male mice were purchased from Jackson Laboratory and injected i.v. once with 5 µl/g saline (naive) or sheep anti-rat glomeruli serum (NTS; Probetex) on day 0, as previously described^18,19^. LL/2 and MC38 cancer cells (5×10^5^) were implanted subcutaneously into the flanks as previously described^11^. Tumor volume was measured with a caliper, and the tumor volume was calculated with the following formula: Width^2^×Length×0.5. An experimental pulmonary metastasis assay was carried out with LLC or MC38 cancer cells (2×10^5^ cells in 200 μL saline) injected into the tail vein, as previously described^11^. At the endpoint, mice were euthanized, and their kidney, lungs, and tumors were excised and weighed.

### Cell Culture

The LL/2 and the MC38 cell lines were purchased from the American Type Culture Collection (ATCC). Cell lines were tested and found to be free of mycoplasma contamination. Cells were cultured in DMEM supplemented with 10% (vol/vol) FBS, 1% streptomycin and penicillin, 1% L-glutamine, and 1% sodium (full medium) pyruvate at 37°C in a humidified atmosphere containing 5% CO2.

### Blood Serum

Blood was obtained from the facial vein with a 4-μm sterile Goldenrod Animal Lancet (MEDIpoint, Inc.). Blood was collected and allowed to clot at room temperature for 2 hours, followed by 15 minutes of centrifugation at 2000g. The serum was immediately aliquoted and stored at −20°C for future use.

### Quantification of circulating factors

Blood serum was analyzed using MILLIPLEX MAP Mouse Cytokine/Chemokine Magnetic Bead Panel - Premixed 32 Plex - Immunology Multiplex Assay (MCYTMAG-70K-PX32, MilliporeSigma) and Proteome Profiler Mouse XL Cytokine Array (ARY028, R&D Systems Inc,) according to the manufacturer’s instructions. In addition, the quantification of epidermal growth factor (EGF) protein levels in the serum was performed with the Mouse EGF Quantikine ELISA Kit (MEG00, R&D Systems Inc.) according to the manufacturer’s instructions.

### Flow cytometry

Single-cell suspensions of kidney, tumor, spleen, blood, and bone marrow were washed with cell staining buffer twice (300g, 5 min, 4⁰C) and incubated in 100 μl of Cell Staining Buffer (10 ml; 420201, Biolegend) with TruStain FcX™ (1:100; 101320, Biolegend) and True- Stain Monocyte Blocker™ (1:20; 426103, Biolegend) for FC blocking (15 min, 4⁰C); then stained with a mixture of fluorescently labeled anti-mouse antibodies (30 min, 4⁰C; **Supplementary Table 1**) in Cell Staining Buffer. Live-dead cell discrimination was performed using Zombie Aqua™ Fixable Viability Kit (423102, Biolegend) according to manufacturer instructions. Following staining, cells were washed three times (300g, 5 min, 4⁰C) in cell staining buffer and kept in FluoroFix™ Buffer (422101, Biolegend) before run in a BD FACSymphony™ A5 Cell Analyzer (BD Biosciences). Analysis was performed using Cytobank (Cytobank.org) or FlowJo™ v10.8 Software (BD Life Sciences).

### Hematoxylin and Eosin Staining

Tumors were fixed in 4% formaldehyde overnight, embedded in paraffin, subsequently serially sectioned at 6-μm intervals, and then mounted on slides. Hematoxylin and eosin staining was performed according to the standard protocol.

### Immunofluorescence Staining

Tumors were fixed in 4% formaldehyde overnight, embedded in optimal cutting temperature compound, and serially sectioned at 10-μm intervals. Frozen tumor sections were stained for CD31 (BD Biosciences, 553370) and Ki67 (Abcam, ab16667) and counterstained with DAPI, as previously described^11^. Each section was fully scanned; for each analysis, 5 fields were randomly chosen and blindly and automatically analyzed with ImageJ software^20^. For every dot plot of image analysis, each dot represents the mean of the values taken from 5 fields, derived from a single mouse.

### RNA Extraction and BulK RNA-Seq

Total RNA was extracted from bone marrow with RNeasy Plus Micro Kit (No. 74034, Qiagen) according to the manufacturer’s instructions. Total RNA was quantified with Qubit RNA HS Assay Kit (Thermo Fisher Scientific) and quality was assessed using RNA ScreenTape on 4200 TapeStation (Agilent Technologies). For sequencing library generation, the Truseq Stranded mRNA kit (Illumina) was used with an input of 100 nanograms of total RNA. Libraries were quantified with Qubit dsDNA HS Assay Kit (Thermo Fisher Scientific) and the average library size was determined using D1000 ScreenTape on 4200 TapeStation (Agilent Technologies). Libraries were pooled and sequenced on NovaSeq 6000 (Illumina) to generate 30 million single-end 50-base pair reads for each sample. Partek Flow software (version 2.7) was used for data analysis.

### Single-cell RNA sequencing (scRNA-Seq) and analysis

Sample processing for scRNA-seq was done using Chromium Next GEM Single Cell 3’ Kit v3.1 (1000269, 10X Genomics) according to the manufacturer’s instructions. Cell concentration was used to calculate the volume of single-cell suspension needed in the reverse transcription master mix, aiming to achieve approximately 8,000 cells per sample. cDNA and libraries were prepared according to the manufacturer’s instructions (10X Genomics). Sequencing reads were processed through 10X Genomics Cell Ranger (v.3.1.0) and aligned to GRCm38. Scanpy40 was used for downstream data analysis. Cells with fewer than 400 genes expressed or more than 5,000 genes expressed were filtered out. Genes that were expressed in less than 50 cells were removed. Cells with 10% or more mitochondrial content were also removed. Scrublet41 was used to remove doublets. Harmony42 was used to harmonize the data from the different samples. Clustering analysis was performed using Leiden clustering with a resolution set to 0.8. Initial cluster annotation was performed using SingleR43.

### Statistics

Data are presented as mean±SE. All mice were included in each statistical analysis unless they were euthanized for humane reasons before the experimental endpoint. Experimental groups were blinded to the experimentalists during data collection. Animals were selected for each group in a randomized fashion. The statistical significance of tumor volume was determined by Two-way repeated-measures ANOVA followed by the Bonferroni posttest. The statistical significance of immune cell populations was determined by Two-way ANOVA followed by Sidak multiple comparisons tests. Comparison between several means was analyzed by One-way ANOVA followed by the Tukey posttest. Comparison between 2 means was performed by a 2- tailed Student t-test. Analyses were performed with GraphPad Prism 9 software (La Jolla, CA). Values of P<0.05 were accepted as statistically significant.

## Results

### Glomerular immune injury attenuates tumor growth

A single injection of nephrotoxic serum (NTS; anti-glomerular IgG) can cause an acute inflammatory reaction in the kidney leading to glomerular injury^21^. Here, we studied whether glomerular immune injury affects non-renal tissue pathology. We used the Nephrotoxic Nephritis (NTN) mouse model to induce glomerular immune injury in C57Bl/6 mice. Saline-injected mice (naive) served as the control group (Figure 1A). Two weeks after saline or NTS injection, we randomly split each group into two groups and challenged the mice with a xenograft lung cancer model. NTS-injected and naive mice were implanted subcutaneously into the flank with LL/2 cells or saline. Tumor growth was monitored over time until the experiment reached a humane endpoint (Figure 1A). Mice injected with only saline (naive) or NTS did not develop tumors. We detected decreased tumor volume in NTS+ LL/2-implanted mice compared with tumors implanted in control mice (Figure 1B). Consistent with tumor volume measurements, the tumor weight of NTS+ LL/2-implanted mice were lower than the tumor weight of the control mice (Figure 1C). We did not detect lung metastatic lesions at the endpoint in this experimental design.

**Figure 1:**
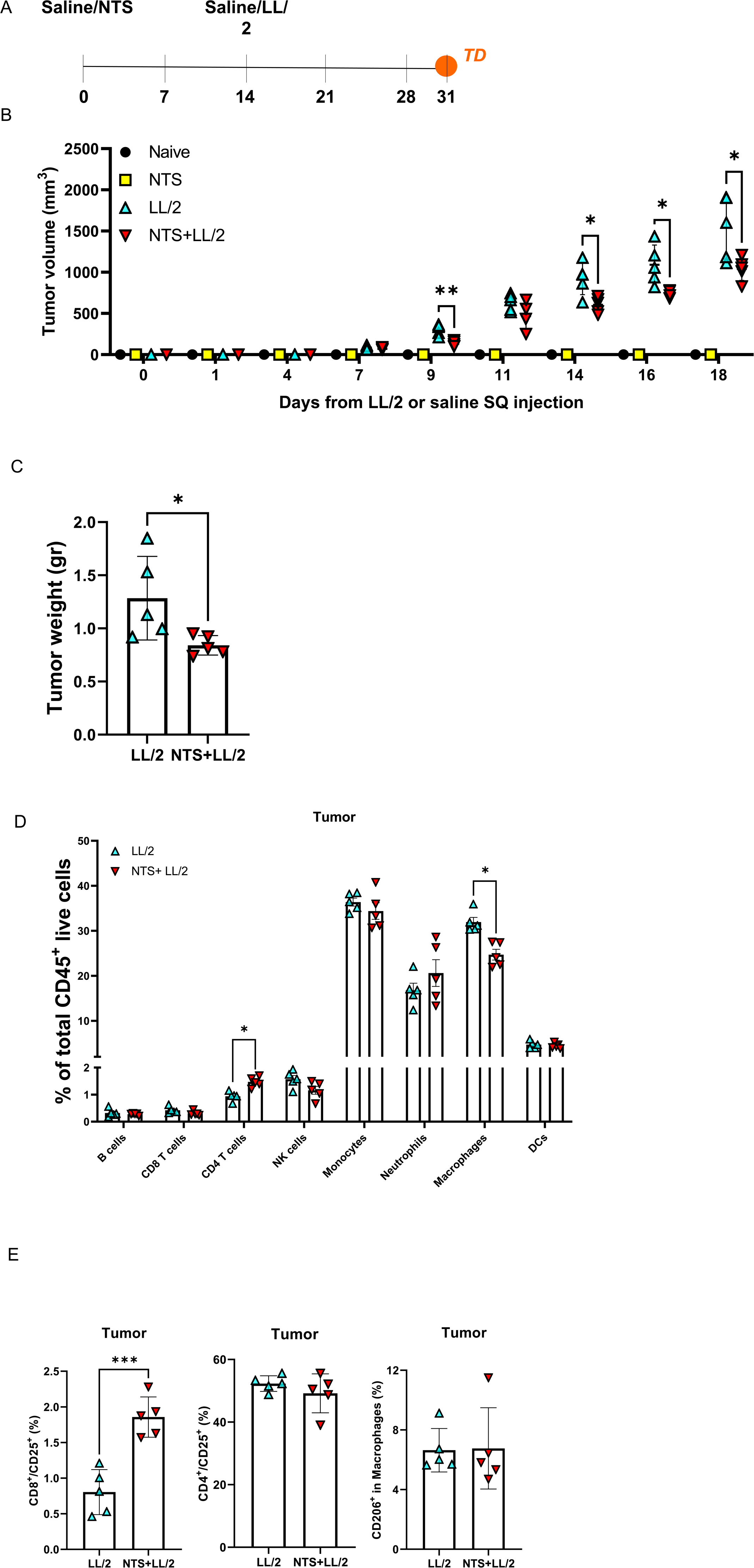
Glomerular immune injury decreases tumor growth. **A**, A schematic diagram of the Lewis lung carcinoma (LL/2) model. Naïve (n=5), NTS-injected (n=5), saline + LL/2 (n=5), or NTS +LL/2 (n=5). Mice were subcutaneously implanted into the flanks with saline or LL/2 cells (0.5×10^6^ cells per mouse) two weeks after the saline or NTS- injection. **B**, Tumor volume was monitored over time using the formula Width^2^×Length×0.5. **C**, Tumor weight at endpoint. Each dot represents 1 mouse. **D**, Single-cell suspensions of tumor cells were stained to identify major immune cell populations and analyzed by flow cytometry. Percentage of B cells (CD45^+^/CD19^+^), CD4 T cells (CD45^+^/CD4^+^), CD8 T cells (CD45^+^/CD8^+^), NK cells (CD45^+^/NK1.1^+^), Monocytes (CD45^+^/CD11b^+^/Ly6C^high^), Neutrophils (CD45^+^/CD11b^+^/Gr-1^+^/Ly6C^Low^), Macrophages (CD45^+^/CD11b^+^/F480^+^) and Dendritic cells (CD45^+^/MHC-II^+^/ F480^-^/CD11c^+^) of total CD45^+^ cells is showed. In addition, cells were stained for CD25, a T-cell activation marker, and CD206 the M2 macrophage marker. **E**, Percentage of activated CD8 T cells (CD8^+^/CD25^+^), activated CD4 T cells (CD4^+^/CD25^+^) of total CD4 or CD8 T cells respectively, and the percentage of M2 macrophages of total macrophages. Data are presented as mean±SD. Two-way repeated-measures ANOVA followed by Bonferroni posttests (**B**), Two-way ANOVA followed by Sidak multiple comparisons tests (**D**), or Student t-test (**C** and **E**). *P<0.05, **P<0.01, ***P<0.001.

To determine whether the anti-tumor phenotype found in NTS+ LL/2-implanted mice was the result of angiogenesis or increased cell proliferation, we immunostained tumor sections with anti-CD31 (endothelial cell marker) and Ki67 (cell proliferation marker). This analysis revealed no substantial differences in the number of blood micro vessels per field between tumor sections of both groups (Figure S1A-B). In addition, there was no statistically significant difference between Ki67 staining in tumor sections of both groups (Figure S1A, C). Moreover, tumor sections stained with H&E showed similar histologic features including necrosis (Figure S1D).

Next, we examined whether blood-circulating factors derived from NS-injected mice could explain the anti-tumor phenotype. We screened serum from the four groups using Luminex (32 cytokines) and the Mouse XL Cytokine Array (111 cytokines). This screening revealed no substantial differences in the concentration of circulating factors (Supplementary Table 1) in the blood of NTS-injected and control mice implanted with LL/2 cancer cells (Figure S2), except for epidermal growth factor (EGF). ELISA confirmed that NTS+ LL/2-implanted mice had less EGF in the blood serum compared with control-LL/2-implanted mice (Figure S3).

To test whether NTN might have systemic effects on immune cell composition in the blood, we used flow cytometry to analyze immune cells from the blood, spleen, kidney, and tumor. We did not detect any significant changes in the percentages of major immune cell populations in the blood or spleen of tumor-bearing mice (control-LL/2 vs. NTS+ LL/2; Figure S4 A-B). In the kidneys of NTS+ LL/2 tumor-bearing mice, we found more CD4 T cells, NK cells, and DC cells compared with control-LL/2 tumor-bearing mice (Figure S4C). Next, we analyzed the immune cells in the tumor. Here, we found more CD4 T cells and fewer macrophages in the NTS+ LL/2 tumors compared with the control tumors (Figure 1D). Although the percentage of CD8 T cells in tumors from both groups were similar, there were higher percentages of activated CD8 T cells (CD25^+^) in the NTS+ LL/2 tumors compared with the control tumors (Figure 1E). These results are consistent with an enhanced immune response in slower-growing tumors.

### The anti-tumor phenotype following NTN is conserved in inflamed tumors

The NTN mouse model has two distinct stages, an acute inflammatory stage between the NTS injection until three weeks and a chronic stage from three weeks onwards ^22^. To gain further insights into the connection between glomerular immune injury and tumor growth at different stages of NTN, we implanted LL/2 cancer cells 1 week, 6 weeks, or 11 weeks post NTS injection. Consistently, tumors of NTS+ LL/2 implanted 1 week after NTS injection were smaller than those implanted in control mice; they had a higher percentage of CD4 T cells, and more activated CD8 T cells (Figure S5). However, after kidney inflammation was resolved, the tumors of NTS+ LL/2 implanted 6 weeks or 11 weeks post NTS injection were the same size as tumors of control+ LL/2 implanted 6 weeks or 11 weeks post saline injection (Figure 2A-C, F-H) suggesting that the effect of NTS injection on tumor growth is transient. We did not detect any significant changes in the percentage of major immune cell populations or T-cell activation in the tumor of both groups at these time points (Figure 2A-C, F-H, S6A-B).

**Figure 2:**
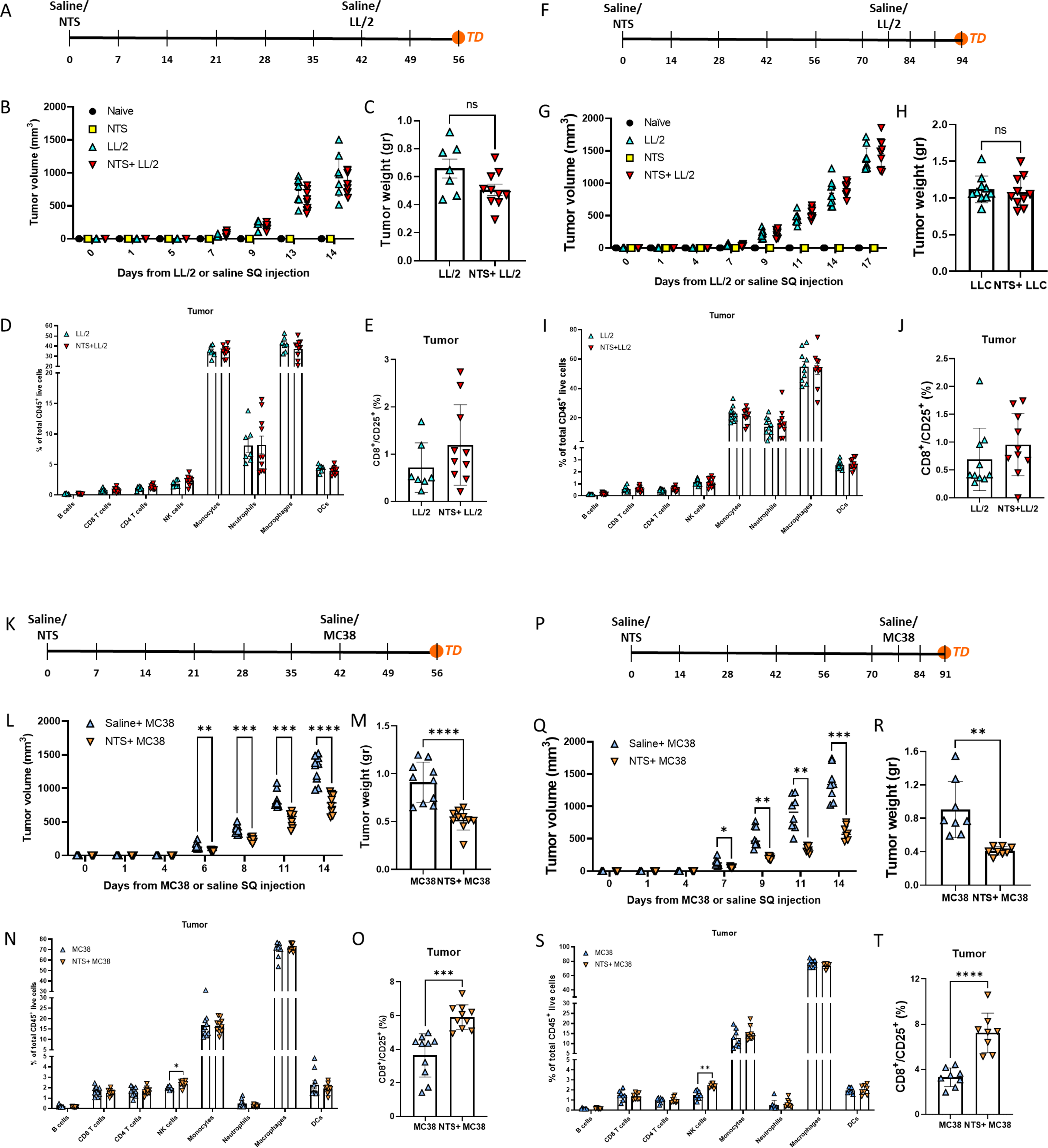
Glomerular immune injury decreases MC38 tumor growth 11 weeks after NTS injection. **A**, A schematic diagram of the Lewis lung carcinoma (LL/2) model. Naïve (n=5), NTS- injected (n=5), saline + LL/2 (n=7), or NTS +LL/2 (n=10). Mice were subcutaneously implanted into the flanks with saline or LL/2 cells (0.5×10^6^ cells per mouse) six weeks after the saline or NTS- injection. **B**, Tumor volume was monitored over time and analyzed as described in **Figure 1B**. **C**, Tumor weight at endpoint. Each dot represents 1 mouse. **D**, Single-cell suspensions of tumor cells were stained and analyzed by flow cytometry as described in **Figure 1D**. Percentage of B cells, CD4 T cells, CD8 T cells, NK cells, Monocytes, Neutrophils, Macrophages, and Dendritic cells of total CD45^+^ cells. **E**, Percentage of activated CD8 T cells (CD8^+^/CD25^+^) of total CD8 T cells. **F**, A schematic diagram of the Lewis lung carcinoma (LL/2) model. Naïve (n=5), NTS-injected (n=5), saline + LL/2 (n=10), or NTS +LL/2 (n=10). Mice were subcutaneously implanted into the flanks with saline or LL/2 cells eleven weeks after the saline or NTS- injection. **G**, Tumor volume was monitored over time and analyzed as described in **Figure 1B**. **H**, Tumor weight at the endpoint. Each dot represents 1 mouse. **I**, Single-cell suspensions of tumor cells were stained and analyzed by flow cytometry as described in **Figure 1D**. Percentage of B cells, CD4 T cells, CD8 T cells, NK cells, Monocytes, Neutrophils, Macrophages, and Dendritic cells of total CD45^+^ cells. **J**, Percentage of activated CD8 T cells of total CD8 T cells. **K**, A schematic diagram of the colorectal carcinoma (MC38) model. Control - saline + MC38 (n=10), or NTS + MC38 (n=10). Mice were subcutaneously implanted into the flanks with saline or MC38 cells (0.1×10^6^ cells per mouse) six weeks after the saline or NTS- injection. **L**, Tumor volume was monitored over time and analyzed as described in **Figure 1B**. **M**, Tumor weight at the endpoint. Each dot represents 1 mouse. **N**, Single-cell suspensions of tumor cells were stained and analyzed by flow cytometry as described in **Figure 1D**. Percentage of B cells, CD4 T cells, CD8 T cells, NK cells, Monocytes, Neutrophils, Macrophages, and Dendritic cells of total CD45^+^ cells. **O**, Percentage of activated CD8 T cells (CD8^+^/CD25^+^) of total CD8 T cells. **P**, A schematic diagram of the colorectal carcinoma (MC38) model. Control - saline + MC38 (n=8), or NTS +MC38 (n=8). Mice were subcutaneously implanted into the flanks with saline or MC38 cells eleven weeks after the saline or NTS- injection. **Q**, Tumor volume was monitored over time and analyzed as described in **Figure 1B**. **R**, Tumor weight at the endpoint. Each dot represents 1 mouse. **S**, Single-cell suspensions of tumor cells were stained and analyzed by flow cytometry as described in **Figure 1D**. Percentage of B cells, CD4 T cells, CD8 T cells, NK cells, Monocytes, Neutrophils, Macrophages, and Dendritic cells of total CD45^+^ cells. **T**, Percentage of activated CD8 T cells of total CD8 T cells. Data are presented as mean±SD. Two-way repeated-measures ANOVA followed by Bonferroni posttests (**B**, **G**, **L**, **Q**), Two-way ANOVA followed by Sidak multiple comparisons tests (**D**, **I**, **N**, **S**), or Student t-test (**C**, **E**, **H**, **J**, **M**, **O**, **R**, **T**). **P<0.01, **P<0.01, ***P<0.001.

The LL/2 tumor is poorly immunogenic. Thus, we hypothesized that the durability of the effects of glomerular immune injury might be more persistent in the setting of a more immunogenic tumor. MC38 is derived from a colorectal tumor and is highly immunogenic. To test this hypothesis, we induced kidney injury with NTS and then implanted MC38 cancer cells 6 weeks or 11 weeks post NTS injection. MC38 tumors implanted 6 weeks or 11 weeks post NTS injection were smaller than control tumors (Figure 2D-E, I-J). Analysis of the tumors showed that more NK cells and activated CD8 T cells were present in NTS-injected mice compared to controls. (Figure 2N-O, S-T, S6C-D). These results suggest that the effects of NTS injection on the immune system are persistent and long-lasting.

### NTN induces transcriptomic changes in BMCs

Since we did not detect any significant changes in the percentage of major immune cell populations in the blood of tumor-bearing control mice when compared with tumor-bearing NTS+ LL/2 mice or of naïve mice when compared with NTS injected mice using flow cytometry (Figure S4A), we hypothesized that NTN-associated systemic inflammation might induce changes in bone marrow cells (BM). Thus, we injected NTS or saline and harvested BM cells (BMCs) after 6 weeks. BMCs were analyzed using flow cytometry and bulk RNAseq. Analysis of BM myelopoiesis and major immune cell populations by flow cytometry, however, revealed no significant differences between NTS-injected and control mice (Figure 3A-C).

**Figure 3:**
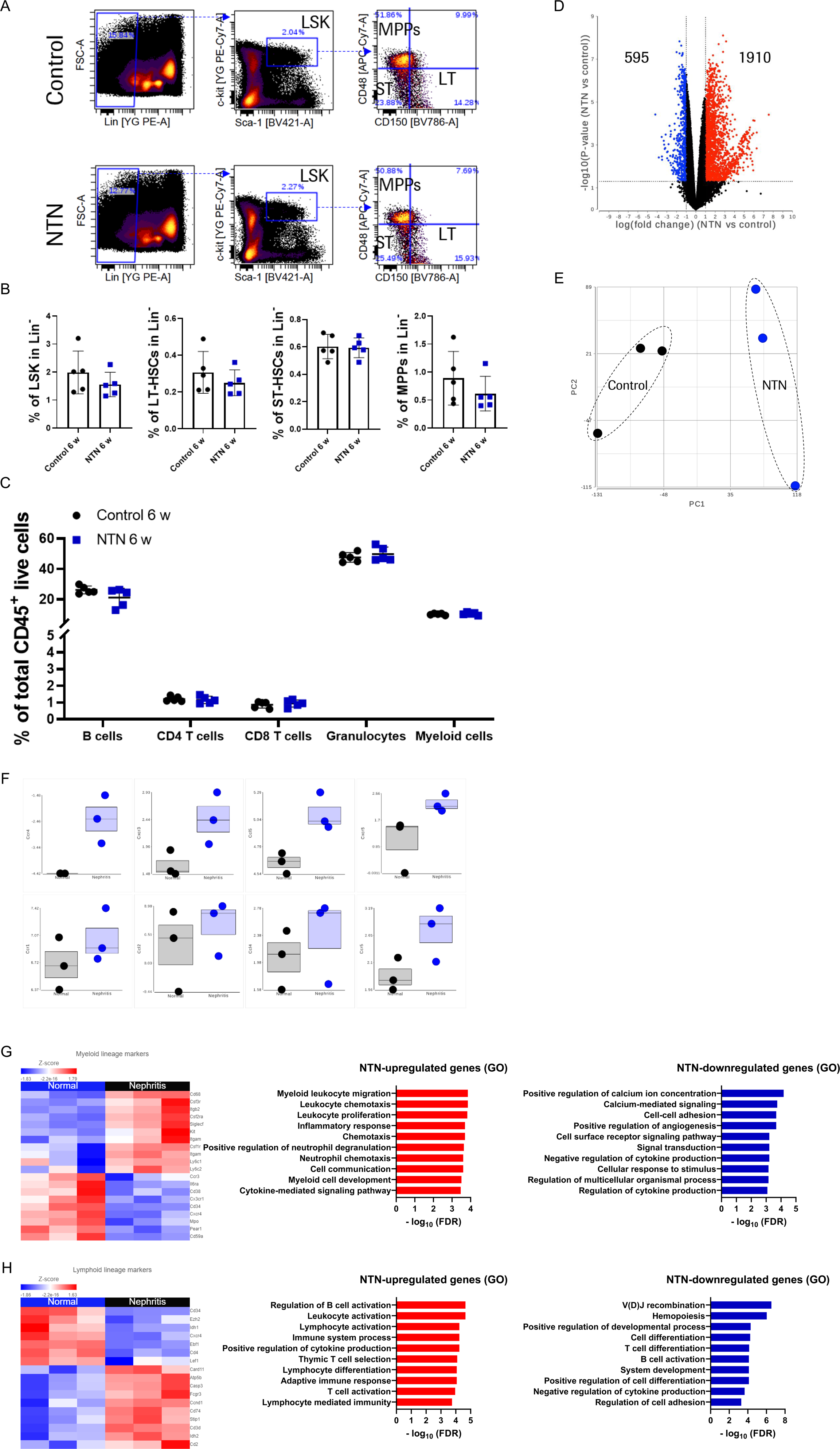
Changes in gene expression of BMCs following NTN. Mice were injected with either saline (control; n=5) or NTS (n=5). Six weeks later, bone marrow cells (BMCs) were harvested and analyzed using FACS. **A**, Representative plots to identify LSK, LT-HSC, ST-HSC, and MPP in total BMCs. **B**, Percentage of LSK, LT-HSC, ST-HSC, and MPP in total BMCs. **C**, Percentage of B cells, CD4 T cells, CD8 T cells, Granulocytes, and Myeloid cells. Each dot represents 1 mouse. Data are presented as mean±SD. Student t-test (**B**) or Two-way ANOVA followed by Sidak multiple comparisons tests (**C**). **D**-**H**, Mice were injected with either saline (control; n=3) or NTS (n=3). Six weeks later, bone marrow cells (BMCs) were harvested and analyzed using RNA-seq. **D**, Differential gene expression of BMCs from NTS-injected mice versus saline-injected controls. The volcano plot shows the adjusted p values distribution and the fold changes. Significant changes are shown in blue (downregulated) or red (upregulated) (FDR < 0.05). **E**, PCA plot of BMC samples from NTS-injected (NTN) and saline-injected (control) mice. **F**, Immune cell homing gene signatures: Ccr4, Cxcr3, Ccl5, Cxcr5, Ccr1, Ccl2, Ccl4, and Ccr5. **G**, Heatmaps of myeloid lineage markers and top overrepresented GO terms including upregulated (red) or downregulated (blue) genes in NTS-injected mice versus saline-injected controls. **H**, Heatmaps of lymphoid lineage markers and top overrepresented GO terms including upregulated (red) or downregulated (blue) genes in NTS-injected mice versus saline-injected controls.

Bulk RNA-Seq analysis of BMCs from NTN-injected and control mice identified 2,505 differentially expressed genes of which 1,910 and 595 genes were significantly upregulated or downregulated, respectively, in BMCs from NTN-injected mice, compared with those from control mice (false discovery rate [FDR] < 0.05) (Figure 3D). Principal component analysis (PCA) showed a clear separation between the two groups (Figure 3E). In addition, BMCs from NTN- injected mice demonstrated strong immune cell homing gene signatures^23^ as shown by upregulation of *Ccr4, Cxcr3, Ccl5, Cxcr5, Ccr1, Ccl2, Ccl4,* and *Ccr5* (Figure 3F). Moreover, we observed alterations in the gene signatures of both myeloid lineage markers and lymphoid lineage markers following NTN (Figure 3G-H). Gene Ontology (GO) enrichment analysis revealed that several GO terms, including “myeloid leukocyte migration”, “inflammatory response”, “positive regulation of neutrophil degranulation”, “neutrophil chemotaxis”, “positive regulation of cytokine production” and activation of B and T cells were overrepresented in the upregulated genes following NTN, whereas terms such as “Calcium-mediated signaling”, “cell-cell adhesion”, or “negative regulation of cytokine production” were overrepresented in the downregulated genes (Figure 3G-H). These findings reveal long-term transcriptomic changes of BMCs that occur after NTS-induced kidney injury.

### Transcriptional alteration in the blood following NTN

To identify whether the transcriptional alternations in the BMCs are reflected in the blood, we analyzed white blood cells (WBCs) of non-tumor-bearing control and NTS-injected (NTN) mice 6 weeks following saline or NTS injection using scRNA-Seq. Two-dimensional Uniform Manifold Approximation and Projection (UMAP) of 10,461 and 5,945 cells in the NTN and control WBCs, respectively, followed by unsupervised clustering analysis identified 14 distinct clusters of cells (Figure 4A). The cell types of each cluster were determined using SingleR using Immgen as the reference (Immgen.org). We observed the same clusters in both groups (Figure 4A). However, a joint UMAP containing both the NTN and control datasets revealed the transcriptional differences mainly in neutrophils and γδ T cells (Figure 4B). Therefore, we used pseudobulk differential expression (DE) analysis to compare γδ T cells (Figure 4C) and neutrophils (Figure 4D, E) of control and NTS-injected mice. The top significantly enriched GO terms in γδ T cells included terms such as “metabolic process” and “regulation of immune system process” (Figure 4F). In neutrophils cluster1 terms such as “lymphocyte chemotaxis” and “chemokine- and cytokine-mediated signaling pathways” (Figure 4G) were overrepresented. In neutrophils cluster 2, metabolism-related terms like “regulation of cellular metabolic process” and “regulation of nitrogen compound metabolic process” were overrepresented in the upregulated genes following NTN (Figure 4H); consistent with the previously described contribution of immuno-metabolic pathways to trained immunity initiation^14^.

**Figure 4:**
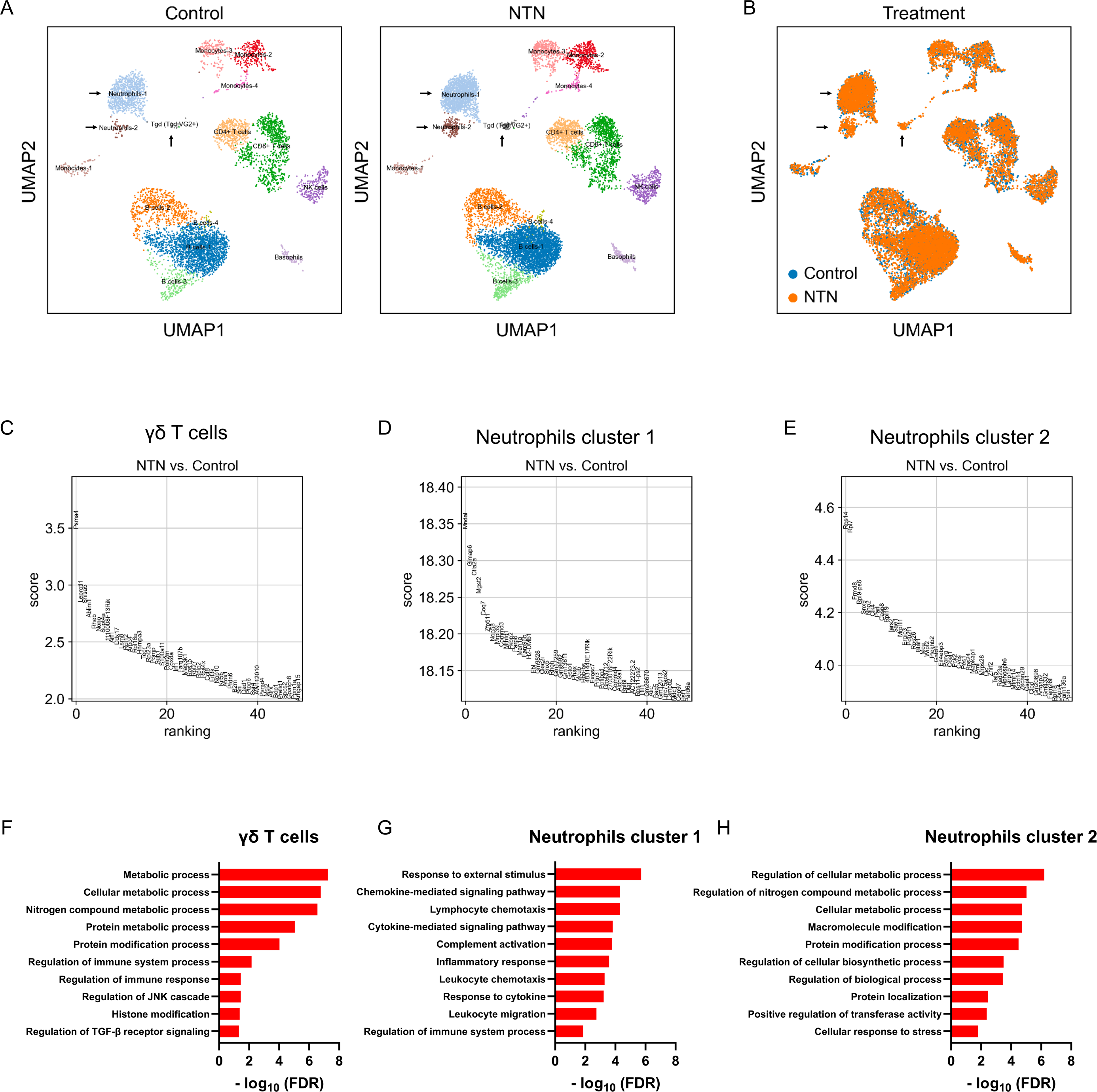
Transcriptional changes in neutrophils. Mice were injected with either saline (control; n=5) or NTS (n=5). White blood cells (WBCs) were harvested, pulled, and analyzed six weeks later using single-cell RNA seq. **A**, Uniform Manifold Approximation and Projection (UMAP) representation of 5945 WBCs of control mice and 10461 WBCs of NTN mice at six weeks from saline or NTS injection. Labels indicate clusters identified by unsupervised clustering analysis. **B**, UMAP of control and NTS-injected mice. **C**-**E**, Pseudobulk differential expression (DE) analysis showing the most differently expressed genes between control and NTS-injected mice in γδ T cells (**C**), neutrophils-1 (**D**), and neutrophils-2 (**E**) clusters. The top 50 upregulated rated genes are shown. **F**-**H**, Upregulated overrepresented GO terms in γδ T cells (**F**), neutrophils-1 (**G**), and neutrophils-2 (**H**) clusters.

## Discussion

A previous study showed that mice pre-treated with a single intraperitoneal (IP) injection of β- glucan, a fungal-derived polysaccharide that stimulates the innate immune system, resulted in growth inhibition of implanted B16-F10 and LLC tumors^14^. Here we showed that similar results could be obtained with mice pre-treated with a nephrotoxic serum that injures the kidneys using two different syngeneic tumors, LLC and MC38. The nephrotoxic serum model of kidney injury that we used involves a sheep antisera prepared against rat glomeruli. This model is an antibody-mediated auto-immune model that resembles the pathogenesis of Goodpasture’s disease or systemic lupus erythematosus. Our studies suggest that multiple methods of immune system stimulation can enhance anti-tumor immunity.

In the previous study, β-glucan likely induces stimulation of myeloid cells that express receptors like Dectin-1 and CR3 that can recognize β-glucan. Sustained changes to myeloid cells, a process coined, trained immunity, enhances subsequent immune reactions to infectious agents as well as to tumors. Sustained changes to myeloid cells are mediated through epigenetic mechanisms. In the NTN model of kidney injury, immune complexes that are formed likely induce complement activation and activate myeloid cells via Fc receptors. While β-glucan injected IP likely spreads systemically, the localization of the NTS likely occurs mainly in the kidney. Thus, we first sought to identify cytokines and growth factors as potential mediators of our phenotype. Our analysis of over 100 cytokines did not reveal any obvious elevated serum cytokines in the blood except for EGF. While the kidney is a major source of systemic EGF potentially impacting the growth of the tumor, our analysis was most consistent with increased immunity as the cause of decreased tumor growth.

Our analysis of changes to bone marrow hematopoiesis by flow cytometry and by bulk RNA-Seq 6 weeks after NTS support sustained changes to bone marrow precursors. Since the bone marrow is constantly seeding the blood with new myeloid cells and because of the complexity of myeloid cell development, we chose to focus on circulating leucocytes using scRNA-Seq.

Treatment-trained granulopoiesis promotes an anti-tumor phenotype in tumor-associated neutrophils (TANs) following β-glucan treatment. The authors identified that ROS production in TANs plays a key role in the anti-tumor effect^14^. The pathogenesis in the NTN model is initiated by anti-glomerular IgGs that damage the glomerular filtration barrier and induce inflammation^22^. Although it has been shown that inflammation can train the immune system to promote anti-tumor activity^14^ the effect of inflammatory stimuli in the kidney on cancer progression has yet to be studied. Indeed, and similar to the transcriptional changes found following β-glucan treatment, immuno-metabolic pathways were overrepresented in NTN-induced alteration in neutrophils in the blood. In addition, although not statistically significant due to their low numbers, we found similar transcriptional changes in γδ T cells. γδ T cells are pivotal in eliminating tumors and cells infected by pathogens due to their powerful cytotoxic, cytolytic, and distinctive immune-modulating abilities^24^. Thus, these transcriptional alterations can account, at least in part, for the anti-tumor activity of NTN.

Our study indicates a link between distinct diseases that involve systemic processes such as circulating proteins. We, therefore, investigated the possible involvement of immune signaling molecules in NTN–dependent tumor growth. We did not detect changes in most of the circulating proteins measured except for EGF. Nevertheless, we may have overlooked some changes due to our limited panel (Supplementary Table 1). Subsequently, we identified EGF to be reduced in the blood serum following NTN. This finding agrees with previous reports showing the reduction in EGF as a predictor of chronic kidney disease (CKD) progression in patients^25^. Interestingly, although EGF-like peptides are overexpressed in most human carcinomas^26^, the reduced levels of EGF following NTN were maintained in tumor-bearing mice (Figure S3) which may contribute to the anti-tumor activity of NTN.

Collectively, our study suggests that glomerular immune injury promotes anti-tumor activity via the reduction in EGF and the transcriptional changes of immune cells. Future studies in pre-clinical models are needed to prove the underline mechanism.

## Supplementary information

**Figure S1: No differences in proliferation, necrosis, or angiogenesis**. **A**, Representative images of LL/2 tumor sections derived from LL/2 (control) and NTS-injected (NTS+LL/2) mice, stained with anti-CD31(endothelial cells; green), anti-Ki67 (proliferating cells; red), and DAPI (nuclei; blue). Magnification 20X. Quantification of endothelial cell staining (CD31+; **B**) or proliferating cells (Ki67+; **C**) per field of tumor sections. Each dot represents the mean of 5 fields taken from one mouse. Data are presented as mean ± SD. Student’s t-test. **D**, Representative images of LL/2 tumor sections derived from LL/2 (control) and NTS-injected (NTS+LL/2) mice, stained with H&E. Scale bar, 200 μm.

**Figure S2: No difference in circulating factors in the blood of tumor-bearing mice**. Protein levels of Interleukin 1 Alpha (IL1*α*; **A**), Granulocyte-Colony Stimulating Factor (G-CSF; **B**), and C-X-C Motif Chemokine Ligand 5 (CXCL5; **C**) in the blood serum of naïve (n=5), NTS- injected (n=5), saline + LL/2 (n=11), or NTS +LL/2 (n=11) mice determined by Luminex. Data from two independent experiments are pooled. Data are presented as mean±SE. One-way ANOVA followed by Tukey posttests. *P<0.05; **P<0.01; ***P<0.001.

**Figure S3: Decreased EGF in the blood of tumor-bearing mice**. Protein expression of 111 cytokines, chemokines, and growth factors in the blood serum of the 4 groups (mix of n=5 per sample) was determined by Proteome Profiler Mouse XL Cytokine array. **A**, The array membrane. Yellow boxes are framing the Epidermal Growth Factor (EGF) antibodies. B, Protein levels of EGF in the blood serum of saline + LL/2 (n=18), or NTS +LL/2 (n=21) mice determined by ELISA. Data from three independent experiments are pooled. Data are presented as mean±SE. Mann-Whitney test. **P<0.01.

**Figure S4: Immune cell populations in the blood, spleen, and kidney**. Single-cell suspensions of white blood cells (**A**), spleen (**B**), and kidney (**C**) cells were stained to identify major immune cell populations and analyzed by flow cytometry. Percentage of B cells (CD45^+^/CD19^+^), CD4 T cells (CD45^+^/CD4^+^), CD8 T cells (CD45^+^/CD8^+^), NK cells (CD45^+^/NK1.1^+^), Monocytes (CD45^+^/CD11b^+^/Ly6C^high^), Neutrophils (CD45^+^/CD11b^+^/Gr-1^+^/Ly6C^Low^), Macrophages (CD45^+^/CD11b^+^/F480^+^) and Dendritic cells (CD45^+^/MHC-II^+^/ F480^-^/CD11c^+^) of total CD45^+^ cells is showed. Data are presented as mean±SE. Two-way ANOVA followed by Tukey posttests. *P<0.05, **P<0.01.

**Figure S5: Reduced LL/2 tumor growth one-week post-NTS injection**. A schematic diagram of the Lewis lung carcinoma (LL/2) model. Naïve (n=5), NTS-injected (n=5), saline + LL/2 (n=7), or NTS +LL/2 (n=9). Mice were subcutaneously implanted into the flanks with saline or LL/2 cells (0.5×10^6^ cells per mouse) one week after the saline or NTS- injection. **B**, Tumor volume was monitored over time using the formula Width^2^×Length×0.5. **C**, Tumor weight at endpoint. Each dot represents 1 mouse. **D**, Single-cell suspensions of tumor cells were stained to identify major immune cell populations and analyzed by flow cytometry. Percentage of B cells, CD4 T cells, CD8 T cells, NK cells, Monocytes, Neutrophils, Macrophages, and Dendritic cells of total CD45^+^ cells is shown. In addition, cells were stained for CD25, a T-cell activation marker, and CD206 the M2 macrophage marker. **E**, Percentage of activated CD8 T cells (CD8^+^/CD25^+^), activated CD4 T cells (CD4^+^/CD25^+^) of total CD4 or CD8 T cells respectively, and the percentage of M2 macrophages of total macrophages. Data are presented as mean±SE. Two-way repeated-measures ANOVA followed by Bonferroni posttests (**B**), Two-way ANOVA followed by Sidak multiple comparisons tests (**D**), or Student t-test (**C** and **E**). *P<0.05, **P<0.01, ***P<0.001.

**Figure S6: CD4 activation and M2 macrophages in LL/2 and MC38 tumors 6- and 11-weeks post NTS injection**. Percentage of activated CD4 T cells (CD4^+^/CD25^+^) of total CD4 and the percentage of M2 macrophages of total macrophages of LL/2 (**A**, **C**) or MC38 (**B**, **D**) tumors implanted 6 weeks (**A**, **B**) or 11 weeks (**C**, **D**) post-NTS-injection. Data are presented as mean±SE. Student t-test.

**Supplementary Table 1:**
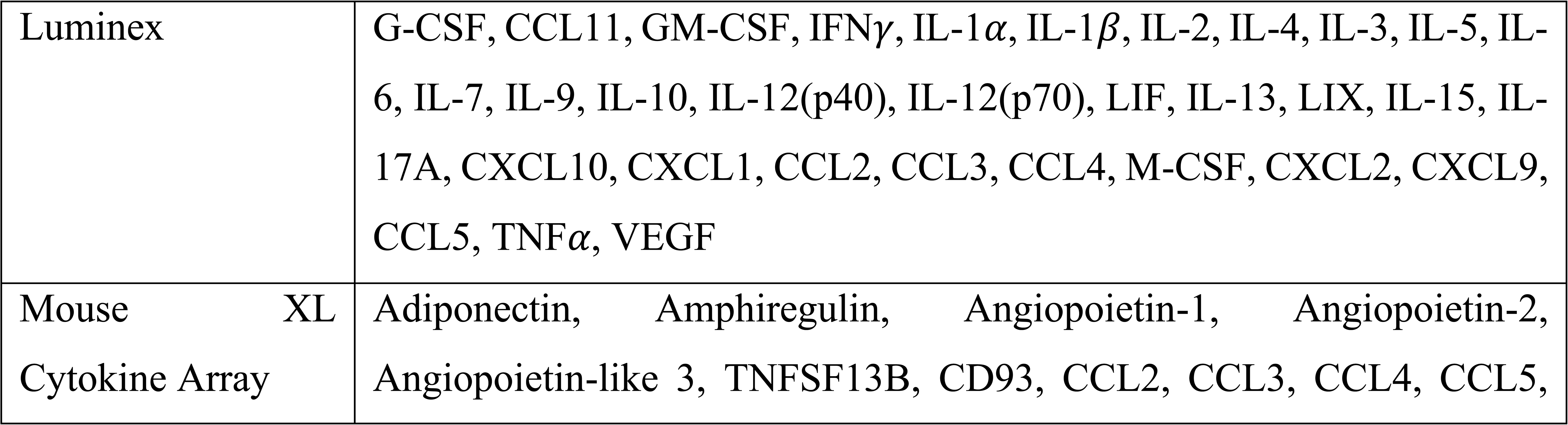
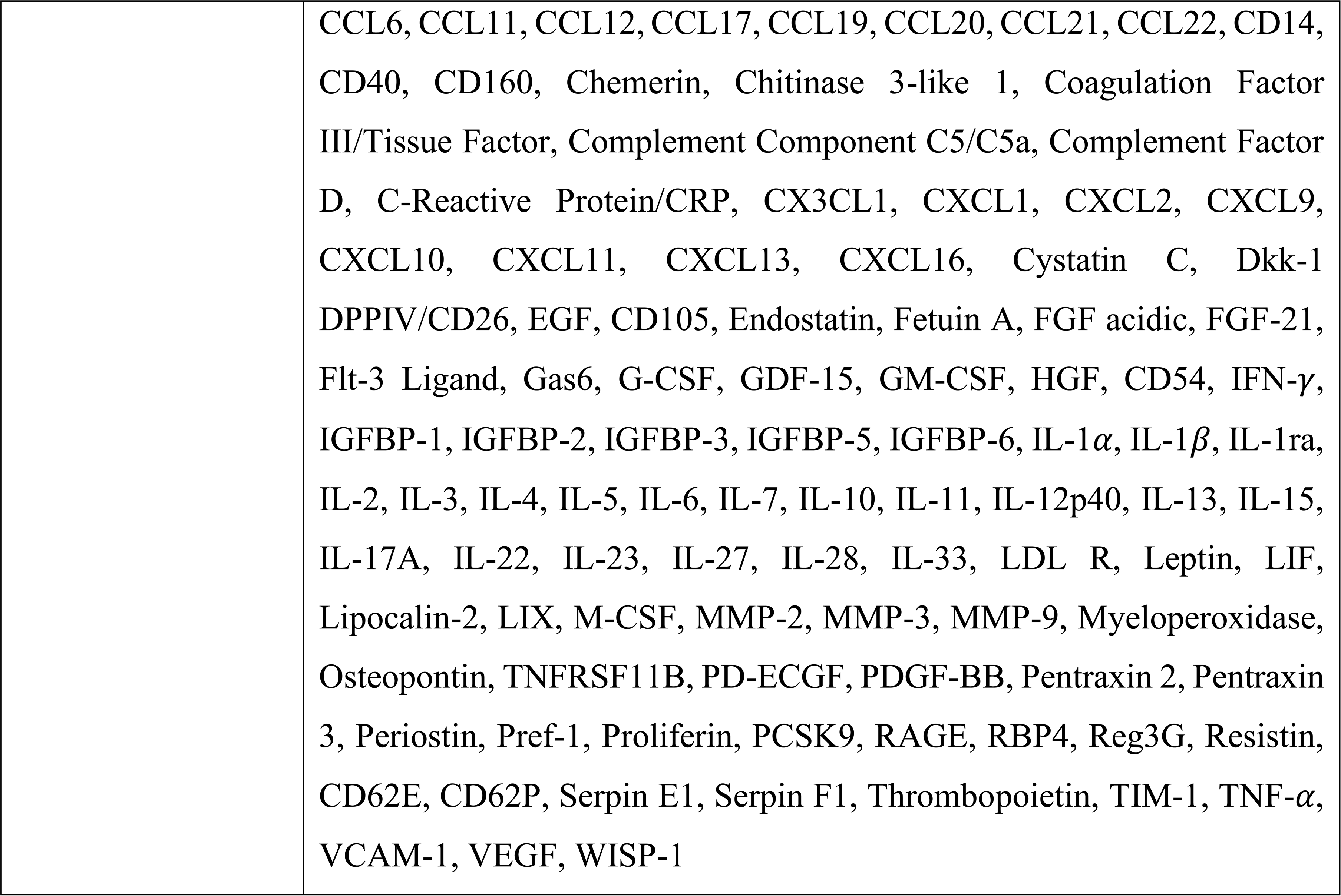
List of circulating factors.

**Supplementary Table 2:** Detailed information on flow cytometry markers used in this study.

| Fluorophore | Target | Manufacture | Catalog # |
| --- | --- | --- | --- |
| PE-Cy7 | CD11c | Biolegend | 117318 |
| PerCP-Cy5.5 | NK1.1 | Biolegend | 156526 |
| FITC | CD62L | Biolegend | 104405 |
| PE | CD8b | Biolegend | 126608 |
| APC-Cy7 | MHC-II | Biolegend | 117324 |
| BV785 | CD45 | Biolegend | 103149 |
| BV711 | CD206 | Biolegend | 141727 |
| BV650 | Ly6G | Biolegend | 127641 |
| BV605 | CD44 | Biolegend | 103047 |
| Zombie Aqua | Live/dead | Biolegend | 423102 |
| BV421 | Ly6C | Biolegend | 128032 |
| BUV737 | CD11b | BD Biosciences | 741722 |
| BUV661 | CD19 | BD Biosciences | 612971 |
| BUV563 | F4/80 | BD Biosciences | 749284 |
| BUV496 | CD4 | BD Biosciences | 741051 |
| BUV395 | CD25 | BD Biosciences | 564022 |
| PE | Lin | Biolegend | 133303 |
| PE-Cy7 | c-Kit | Biolegend | 135111 |
| BV421 | Sca-1 | Biolegend | 108127 |
| APC-Cy7 | CD48 | Biolegend | 103431 |
| BV786 | CD150 | BD Biosciences | 567518 |

## Data availability

The data that support the findings of this study are available upon reasonable request.

## Contributions

S.A. conceived the project, designed and performed experiments, analyzed and interpreted the data, created the figures, and wrote the manuscript. B.K. performed flow cytometry experiments, assisted in data interpretation, and revised the manuscript. J.H.H.L. performed scRNA-seq data analysis. M.C-T assisted in animal experiments. H.M, S.D. Y.L. Z.M. supported bulk RNA-Seq and scRNA-Seq. S.D. performed bulk RNA-seq data analysis. J.D.W. performed pathology analysis and assisted in data interpretation. A.S. oversaw data interpretation and revised the manuscript.

## Ethics declarations Competing interests

This study received funding from Genentech Inc. The funder was not involved in the study design, collection, analysis, interpretation of data, the writing of this article or the decision to submit it for publication. All authors are employees of Genentech Research and Early Development.

## Acknowledgments

We thank our Genentech colleagues in the Research Biology, Laboratory Animal Resources, and Pathology Departments for their support in this study. This work was supported by Genentech.

