## Supplementary Figures for "Glomerular immune injury promotes anti-tumor activity"

**Supplementary Figure 1: No differences in proliferation, necrosis or angiogenesis**

A

LL/2 | NS+ LL/2

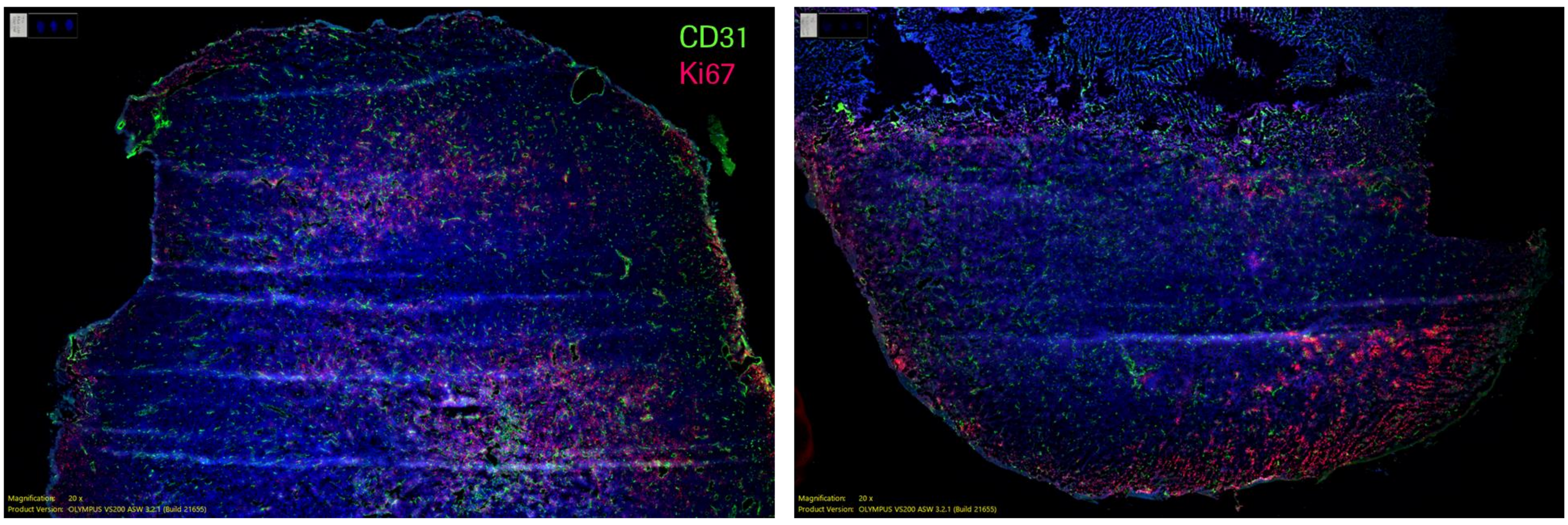


B

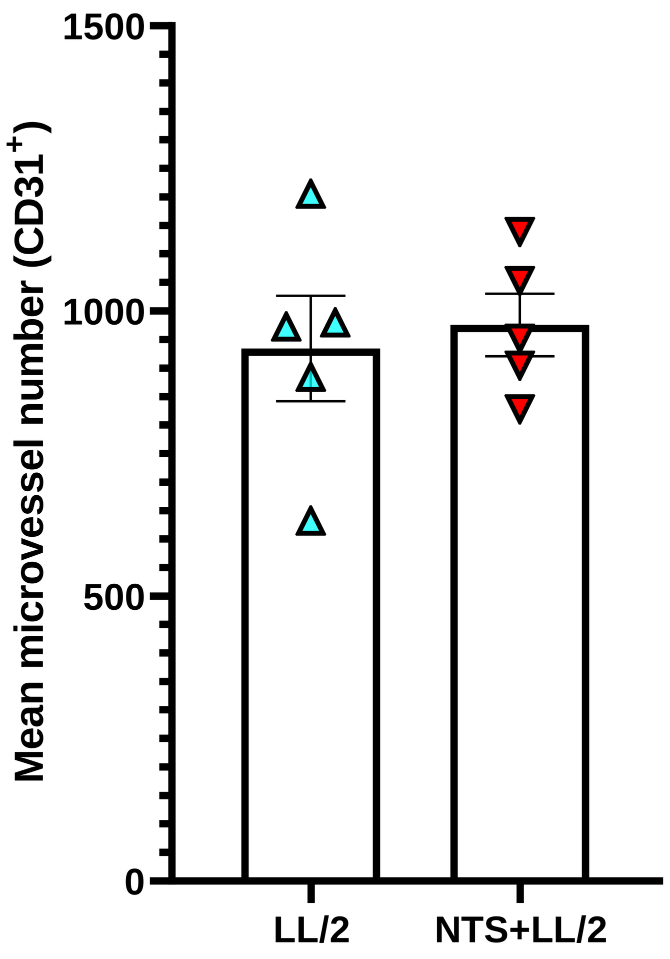


C

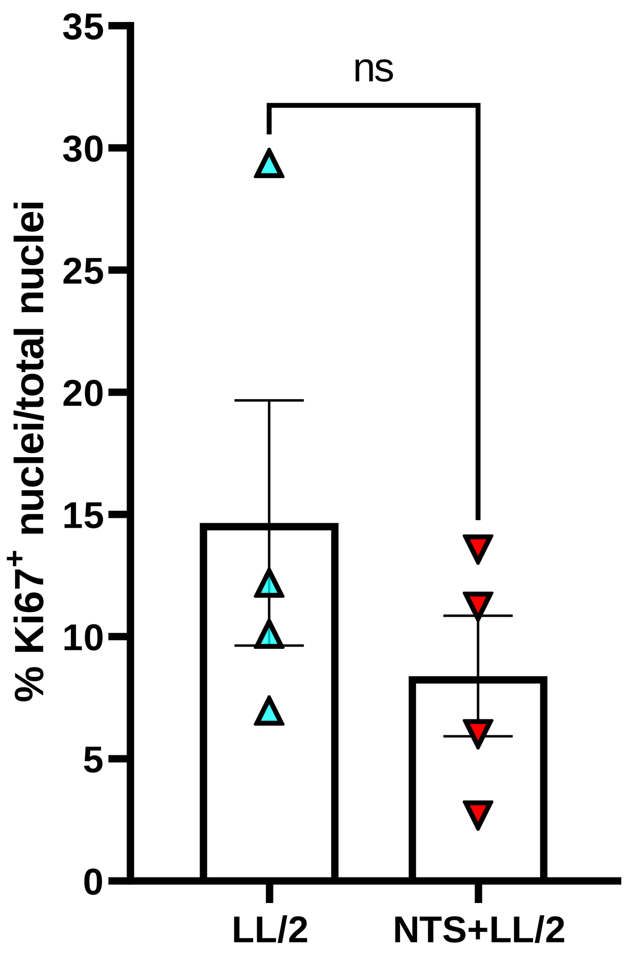


D

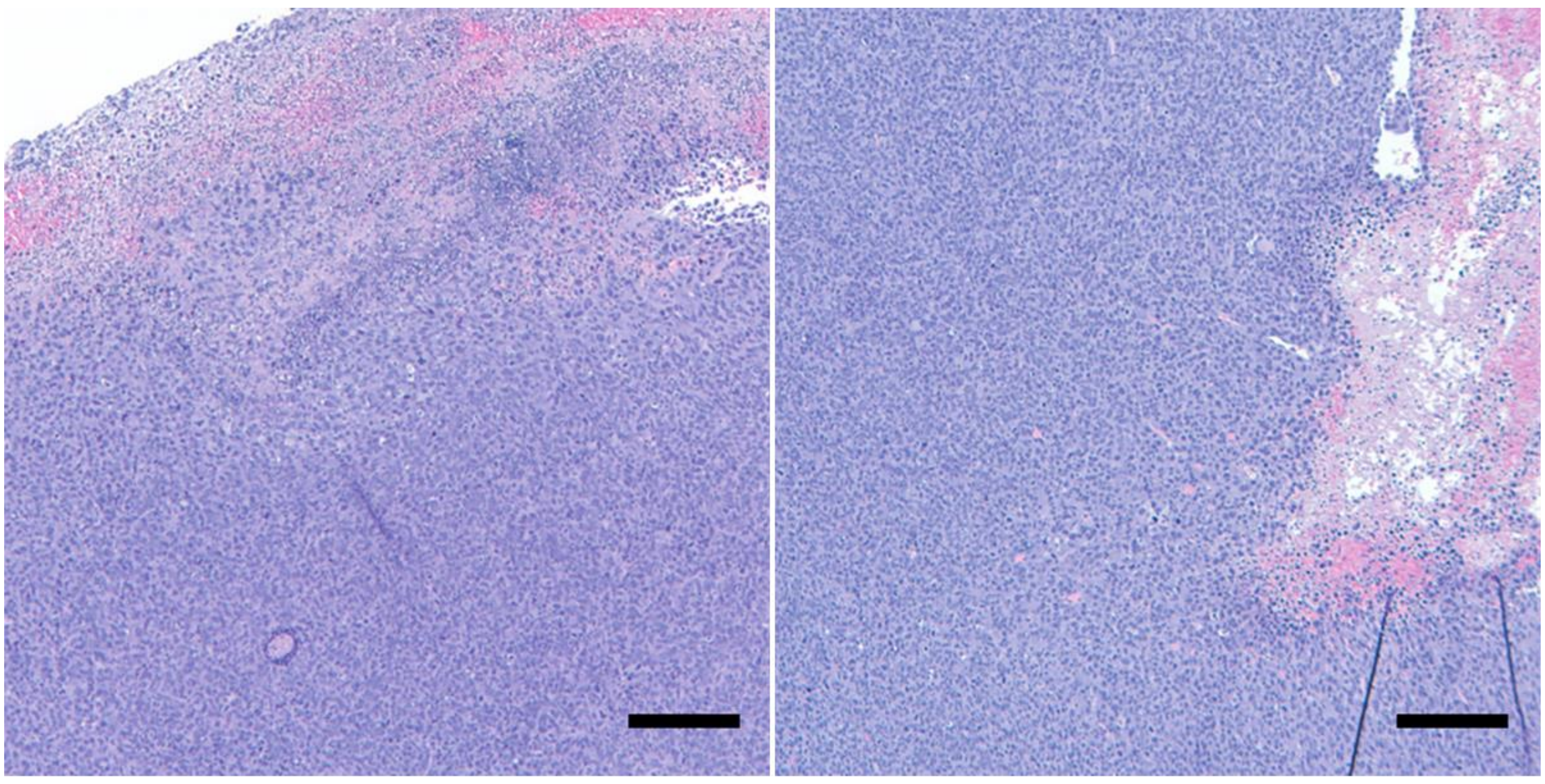

Nephrotoxic Serum, LL/2 | LL/2 Alone

**Supplementary Figure 2: No difference in circulating factors in the blood of tumor-bearing mice**

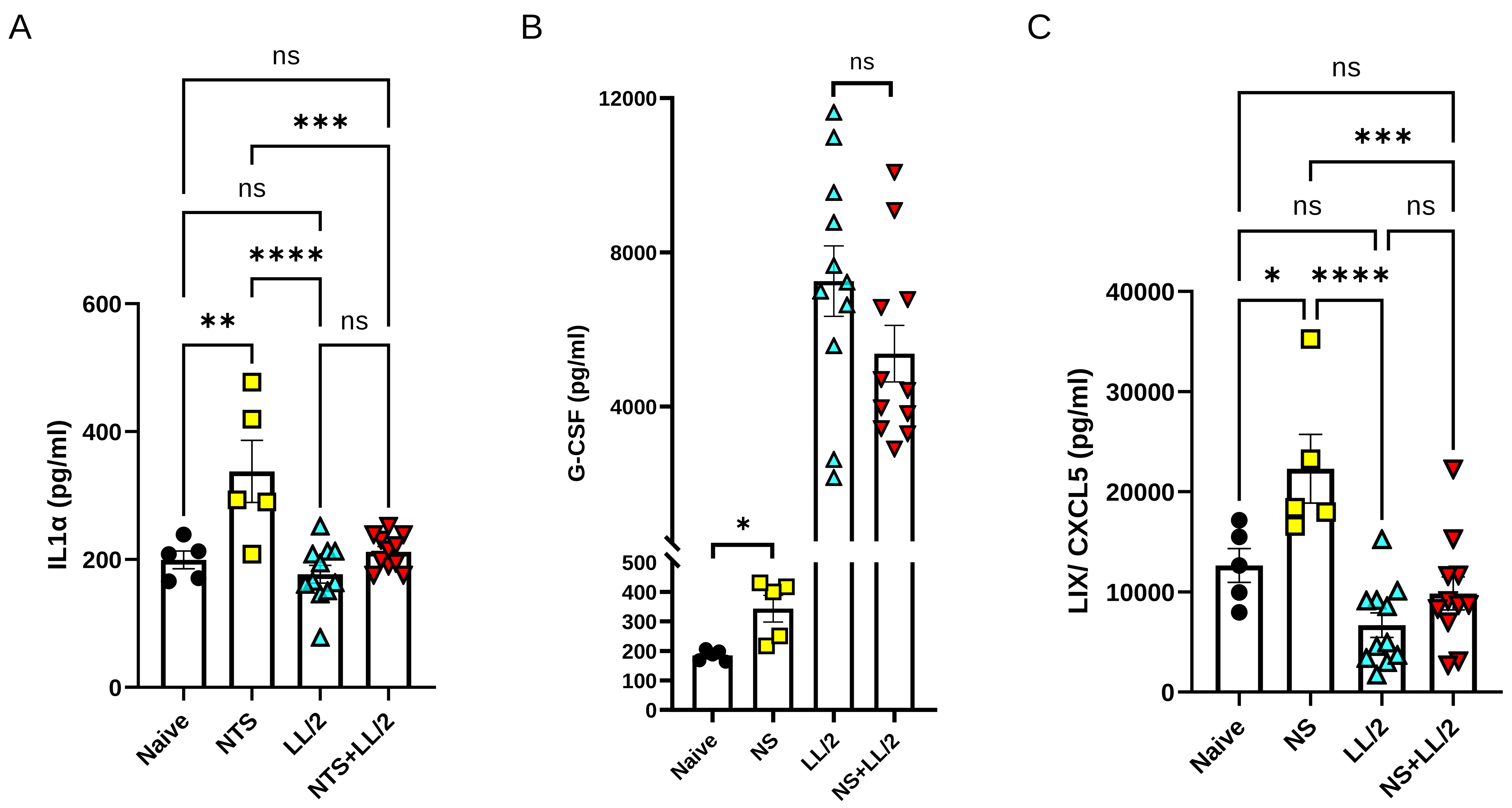

**Supplementary Figure 3: Decreased EGF in the blood of tumor-bearing mice**

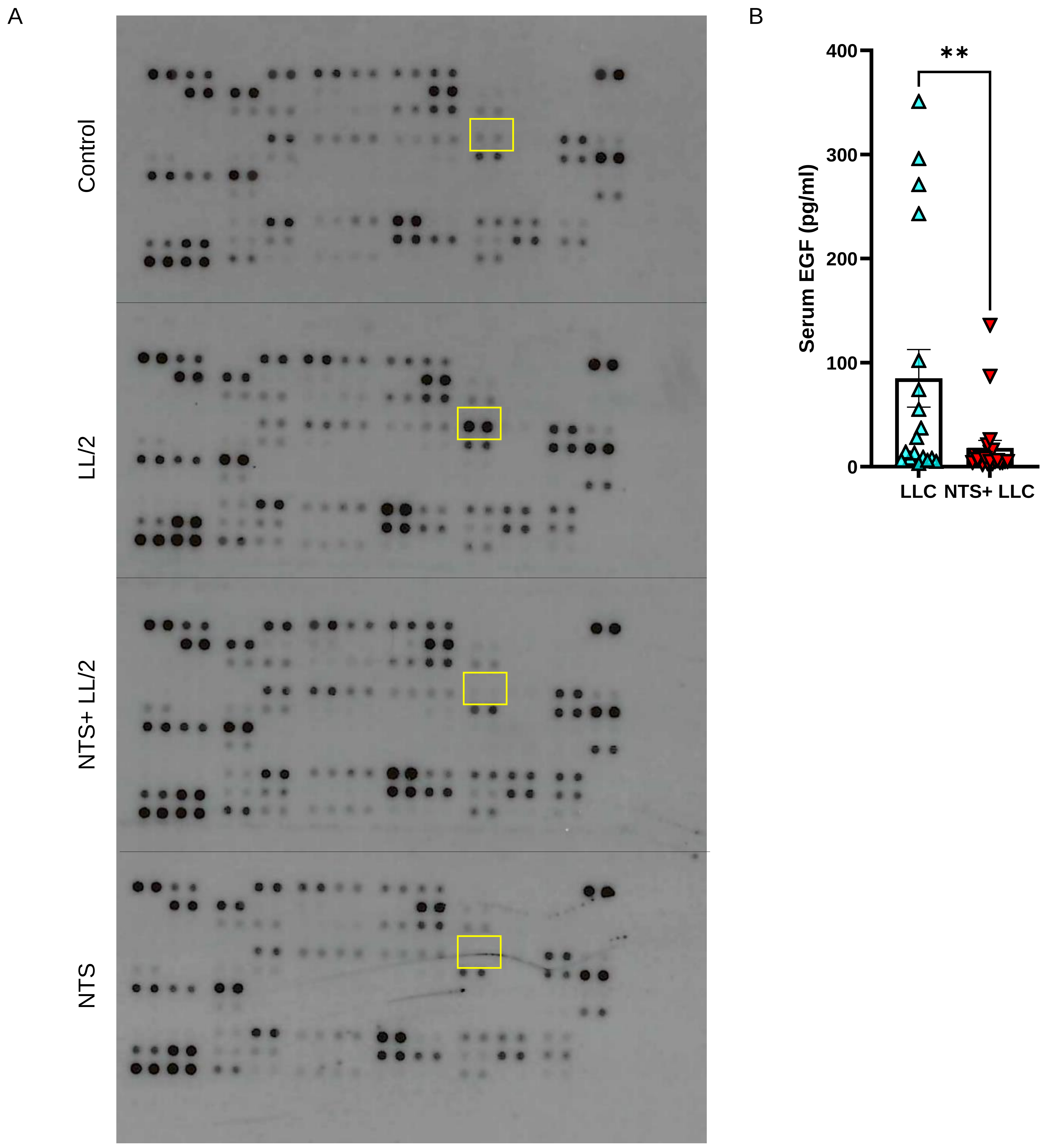

**Supplementary Figure 4: Immune cell populations in the blood, spleen, and kidney**

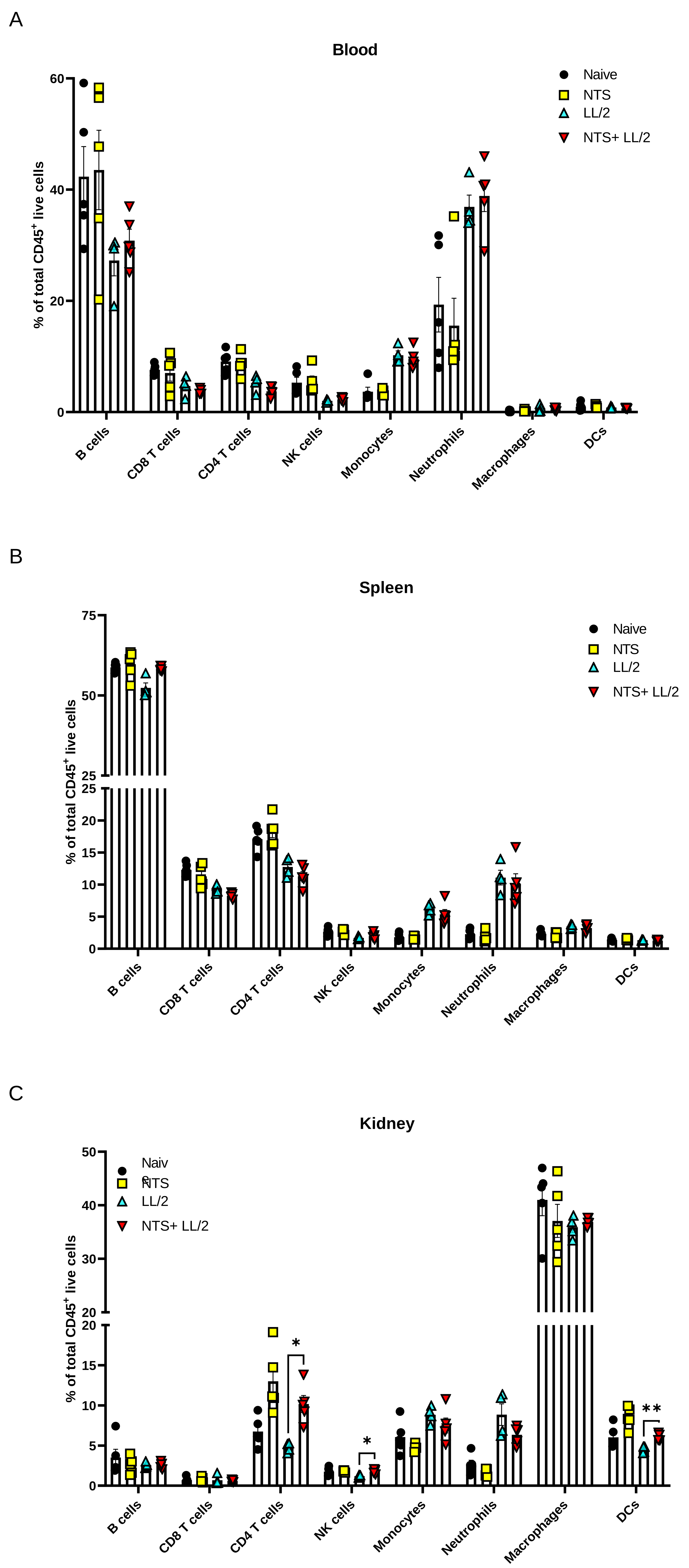

**Supplementary Figure 5: Reduced LL/2 tumor growth one-week post-NTS injection**

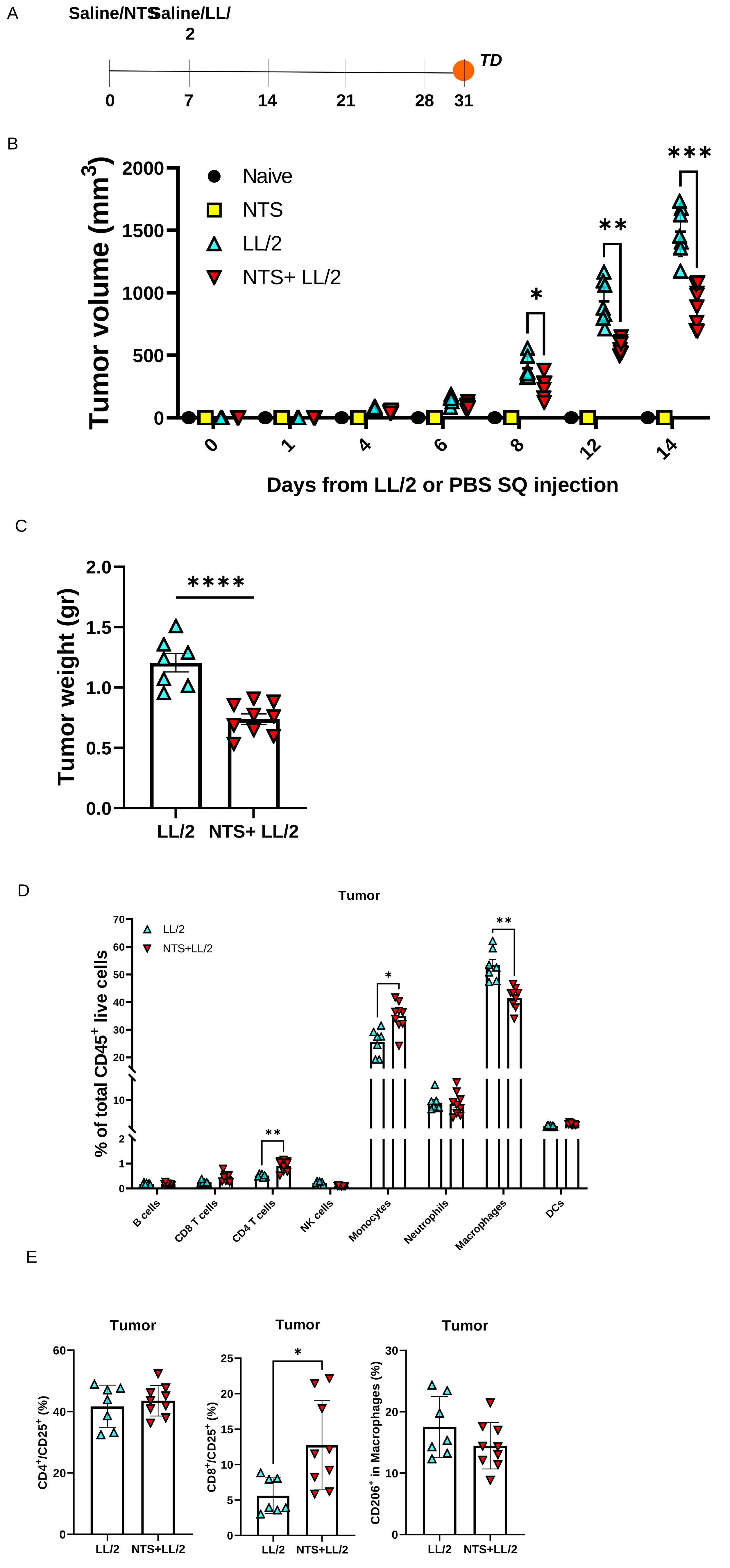

**Supplementary Figure 6: CD4 activation and M2 macrophages in LL/2 and MC38 tumors 6 and 11 weeks post NTS injection**

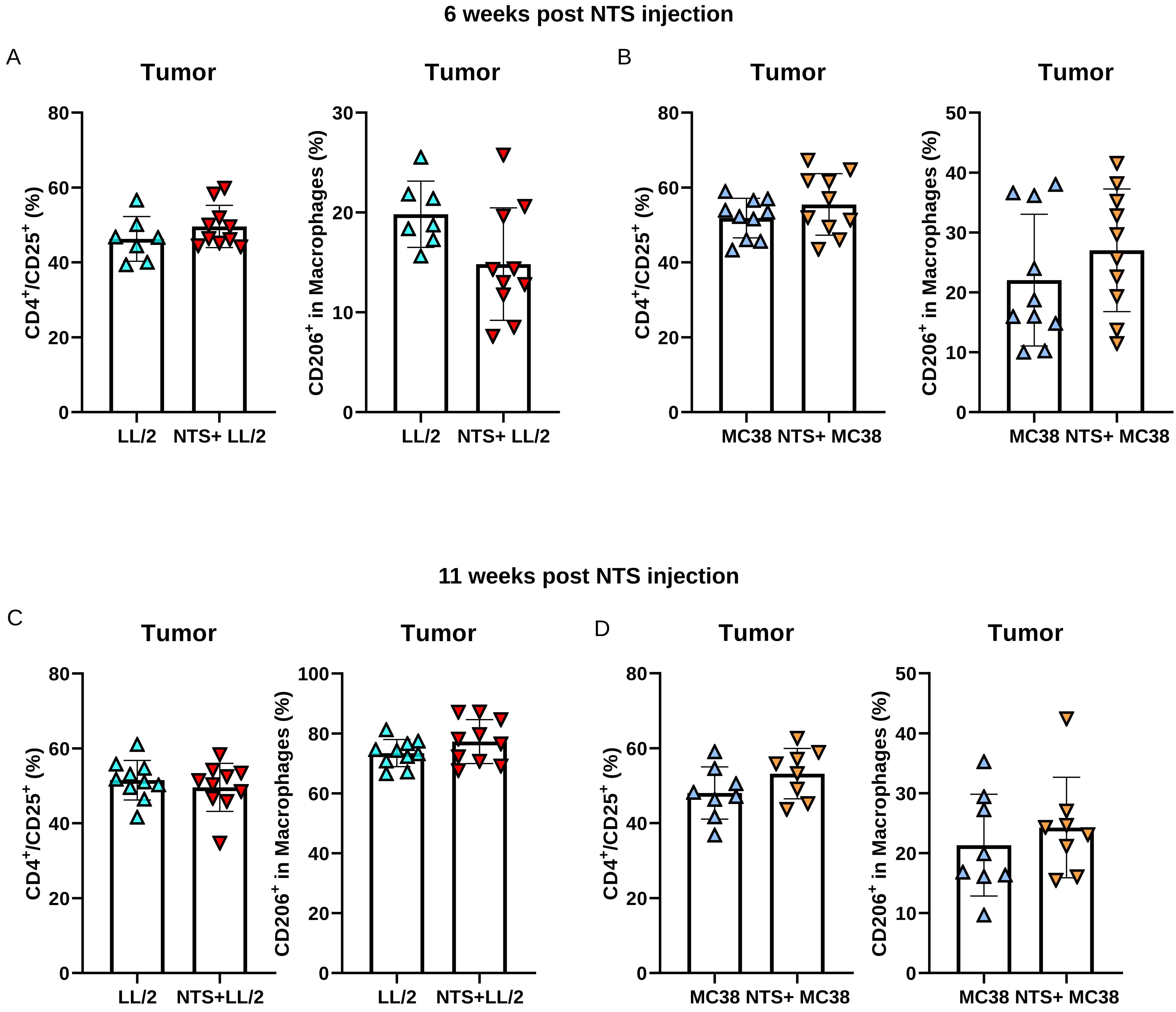
